# Compromised function of the ESCRT pathway promotes endolysosomal escape of tau seeds and propagation of tau aggregation

**DOI:** 10.1101/637785

**Authors:** John J. Chen, Diane L. Nathaniel, Preethi Raghavan, Maxine Nelson, Ruilin Tian, Eric Tse, Jason Y. Hong, Stephanie K. See, Sue-Ann Mok, Marco Y. Hein, Daniel R. Southworth, Lea T. Grinberg, Jason E. Gestwicki, Manuel D. Leonetti, Martin Kampmann

## Abstract

Intercellular propagation of protein aggregation is emerging as a key mechanism in the progression of several neurodegenerative diseases, including Alzheimer’s Disease and frontotemporal dementia. However, we lack a systematic understanding of the cellular pathways controlling prion-like propagation. To uncover such pathways, we performed CRISPR interference (CRISPRi) screens in a human cell-based model of propagation of tau aggregation. Our screens uncovered that knockdown of several components of the ESCRT machinery, including CHMP6, or CHMP2A in combination with CHMP2B (a gene linked to familial fronto-temporal dementia), promote propagation of tau aggregation. We found that knockdown of these genes caused damage to endolysosomal membranes, consistent with a role for the ESCRT pathway in endolysosomal membrane repair. Leakiness of the endolysosomal compartment significantly enhanced prion-like propagation of tau aggregation, likely by making tau seeds more available to pools of cytoplasmic tau. Together, these findings suggest that endolysosomal escape is a critical step in tau propagation.

---

Neurodegenerative diseases are one of the most pressing challenges facing humanity. A formidable roadblock to the development of effective therapies is our incomplete understanding of the underlying molecular and cellular mechanisms. A major breakthrough was the discovery that scrapie, an infectious neurodegenerative disease, is caused by the cell-to-cell propagation of protein aggregates via “prion” forms of the protein (1). In this process, a prion seed converts healthy, native proteins to adopt an aggregated, prion conformer. More recently, findings from numerous, independent studies support the hypothesis that prion-like propagation also underlies common, non-infectious neurodegenerative diseases, such as Alzheimer’s Disease (AD) (recently reviewed in (2)). However, the mechanisms that control aggregate uptake and propagation remain to be fully elucidated, especially in those diseases that involve cytoplasmic proteins. A systematic understanding of these mechanisms is important, both for the development of therapeutics and for furthering our understanding of why specific neuronal subtypes and brain regions are especially susceptible to specific diseases.

Of particular interest to us are the mechanisms controlling propagation of aggregated forms of the protein tau. Tau aggregation is one of the hallmarks of AD and the levels of aggregated tau correlate with cognitive deficits and neuronal loss (3-6). Beyond AD, tau aggregation also defines a number of other neurodegenerative diseases, collectively termed tauopathies, some of which are caused by familial point mutations in tau (7).

Propagation of tau aggregation can be modeled in cultured HEK293 cells that express fluorescently tagged versions of tau, as first established by the Diamond lab (8). In this system, addition of aggregated tau seeds to the culture media causes the fluorescently tagged tau in the cells to convert from a diffuse, soluble form to aggregated puncta. This cell-based model has enabled the characterization of tau species with seeding activity from patient brains (9), and the creation of a minimal synthetic tau that retains seeding capability (10). Furthermore, cell-based models can also be used as a biosensor to detect and propagate distinct prion strains of tau from different tauopathies (11-13). Importantly, seeding of tau aggregation in the cell-based model is predictive of *in vivo* seeding in a mouse model (14).

In addition to their utility as “biosensors” for tau aggregates with prion properties, cell-based models can also be used to elucidate cellular pathways that control propagation of tau aggregation. Previous work from others and us leveraged cell-based models to uncover mechanisms that mediate tau uptake into cells (8, 15-17). In those studies, binding of tau to specific cell-surface heparan sulfate proteoglycans was found to mediate cellular uptake. These results were validated in human iPSC-derived neurons and mouse brain slices (16), supporting the physiological relevance of the cell-based model.

While these studies established the mechanism for tau uptake, the downstream cellular pathways controlling propagation of tau aggregation have not been systematically characterized. We hypothesized that tau aggregation in the cytosol would be influenced by multiple cellular pathways, including those controlling trafficking of tau seeds through the endolysosomal pathway, localization of tau seeds to the cytosol, templated aggregation of soluble tau, and clearance of tau aggregates (**Fig. 1A**).

**Figure 1.**
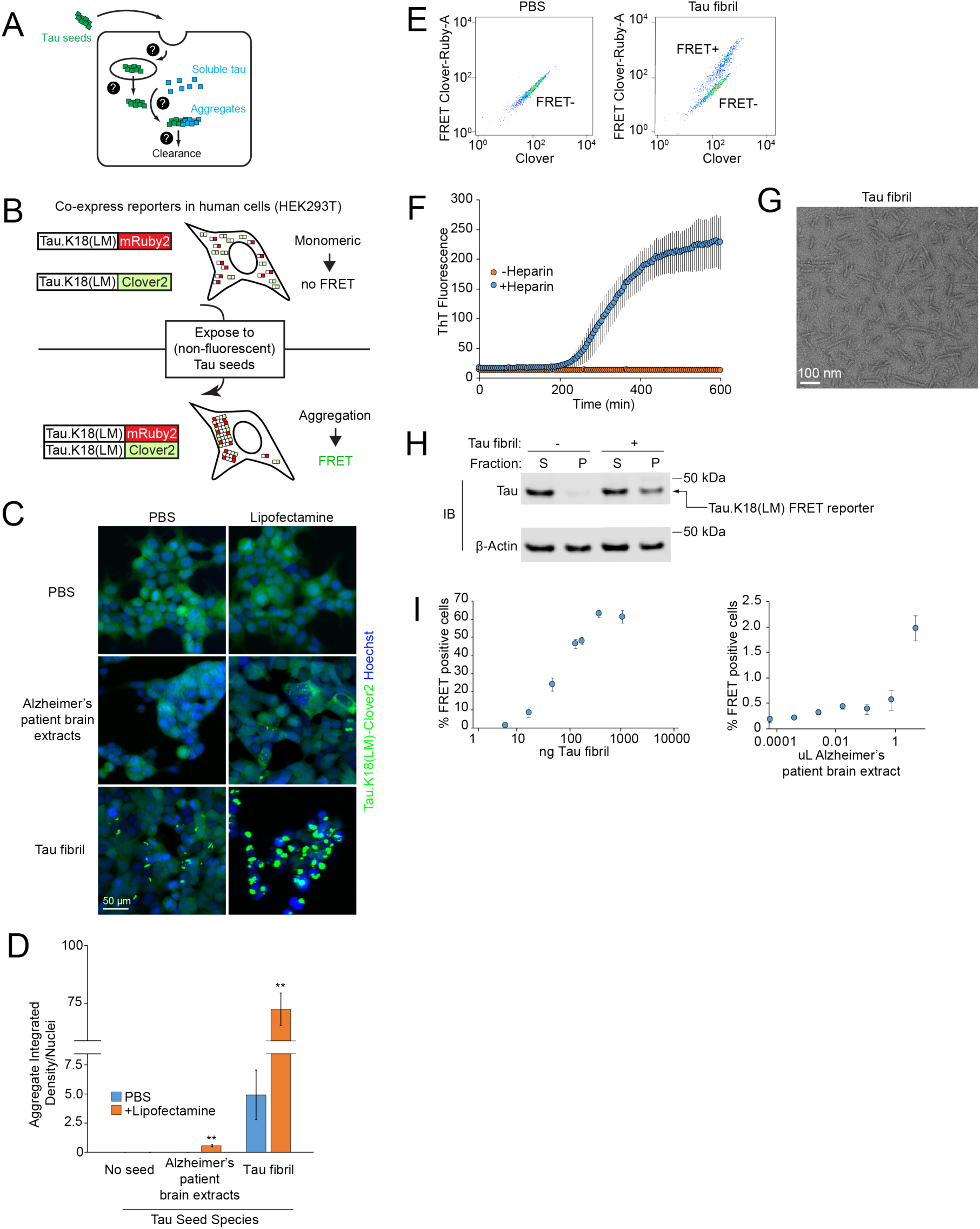
Tau seeds induce tau aggregation in a FRET-based reporter cell line. (**A**) Overview of cellular processes that may control the prion-like tau propagation and aggregation. Question marks represent unknown cellular mechanisms. (**B**) Schematic representation of the FRET-based reporter assay to monitor tau aggregation in HEK293T cells. In the absence of tau seeds, fluorescently labeled tau.K18(LM) is monomeric. Exposure to non-fluorescent tau seeds induces aggregation of the reporter, which can be measured by the formation of tau aggregates by fluorescence microscopy or an increase in FRET intensity by flow cytometry. (**C**) Induction of fluorescent tau aggregates in FRET reporter cell line. Representative images of cells treated with PBS (top row), Alzheimer’s patient brain extracts after 5 days (*second row*), or fibrils of recombinant human 0N4R tau after 2 days (*third row*). For each tau seed, each condition is complexed with (*right column*) or without (*left column*) lipofectamine. Nuclei were counter-stained with Hoechst 33342. (**D**) Comparison of intracellular fluorescent tau aggregates from images in **Fig. 1C**. Integrated density quantification of fluorescent tau aggregates seeded with various tau seeds complexed with *(blue)* or without (*orange*) lipofectamine were quantified and divided by total nuclei per image. n=3 technical replicates (with at least 50 nuclei per image), error bars represent standard deviation, * P<0.05, ** P<0.01 (two-tailed Student’s t test for comparison to PBS (no lipofectamine) control for each tau seeding condition). (**E**) Representative flow cytometry plot of FRET reporter cells after 2 day treatment with PBS (*left*) or tau fibrils (*right*). (**F**) Incubation of recombinant 0N4R tau with heparin and constant agitation at 37° C induces fibrillization. Fibrillization is monitored using an increase in Thioflavin T fluorescence (ex: 440 nm, em: 485 nm), which occurs in the presence (blue) of heparin (10 µg/mL), but not in the absence (orange) of heparin. Error bars represent standard deviation from n=3 technical replicates. (**G**) Representative negative stain electron micrograph of tau fibrils. (**H**) Lysates from FRET reporter cells treated with PBS or tau fibrils for 2 days were fractionated at 1000xg into soluble (S) or pellet (P) fractions, and subjected to SDS-PAGE and immunoblotting using antibodies against tau and β-actin. (**I**) Quantification of % FRET positive cells using flow cytometry across concentration ranges of tau fibrils (*left*) or human Alzheimer’s patient brain extracts (*right*). Error bars represent standard deviation for n=3 technical replicates.

To uncover relevant cellular pathways downstream of tau uptake, we here combine our CRISPR interference-based genetic screening approach (18, 19) with a cell-based model of tau aggregation using fluorescence resonance energy transfer (FRET). Using this approach, we uncover endolysosomal escape of tau seeds as a critical step in the propagation of tau aggregation. Defects in the ESCRT machinery compromise endolysosomal integrity, thereby promoting the escape of tau seeds from endolysosomal compartments and accelerating subsequent templating of tau aggregation in the cytosol. These findings provide insight into the mechanisms of tau trafficking and suggest a source for new potential therapeutic targets.

## RESULTS

### Cell-based model of prion-like propagation of tau aggregation

We established a cell-based model to monitor the prion-like propagation of tau aggregation in HEK293T cells. Such a model had previously been pioneered by the Diamond lab (8) and adapted for flow cytometry using a FRET-based strategy to monitor tau aggregation (20). In this FRET-based strategy, two versions of the tau repeat domain (RD) containing disease-associated P301L and V337M mutations are expressed as fusions with either the FRET donor CFP or the FRET acceptor YFP. When exposed to tau fibrils from recombinant or cell/brain-derived lysate sources, the CFP and YFP tags are brought into close proximity, enabling FRET. We generated a reporter line following a similar strategy (**Fig. 1B**). Instead of the CFP-YFP FRET pair, we used Clover2 and mRuby2, since proteins of this type had been shown to have a very high dynamic range for FRET, with a high Förster radius (21). We selected a monoclonal line for optimal expression and dynamic range of the FRET signal.

In the absence of seeding, these cells showed diffuse intracellular fluorescence without visible aggregates when monitored by fluorescence microscopy (**Fig. 1C,D**) and they appeared as a single population when FRET levels are monitored by flow cytometry (**Fig. 1E**). In contrast, exposure of these cells to extracts from AD patient brains caused the tau reporter constructs to aggregate, as reflected by formation of fluorescent puncta (**Fig. 1C,D**). However, seeded aggregation with brain-derived tau required co-incubation with lipofectamine 2000 (here referred to as lipofectamine) to achieve modest aggregation (**Fig. 1C,D**), consistent with reports from other groups (12-14).

Our goal was to eliminate the use of lipofectamine, because the use of a lipocationic carrier may bypass physiologically relevant uptake or trafficking pathways. Accordingly, we purified monomeric 6xHis-tagged 0N4R human tau from *E. coli* and induced fibrillization with heparin, which we monitored by an increase in thioflavin T fluorescence (**Fig. 1F)** and by negative stain electron microscopy (**Fig. 1G**). We found that treatment of our FRET reporter cells with these tau fibrils caused robust formation of aggregates, even in the absence of lipofectamine. This activity was confirmed using multiple criteria, including formation of puncta by fluorescence microscopy (**Fig. 1C,D**), appearance of a FRET-positive population by flow cytometry (**Fig. 1E**), and biochemical characterization of tau in the insoluble fraction (**Fig. 1H**). Finally, we tested the effects of increasing concentrations of tau fibrils on our FRET-based reporter. We found that 6xHistagged fibrils robustly triggered tau aggregation in a dose-dependent manner across nearly 2 orders of magnitude in concentration in the absence of lipofectamine, as quantified by the percentage of FRET positive cells (**Fig. 1I**). Brain lysates also produced an increase in FRET-positive cells, although the magnitude was more modest. Together, these features make our FRET-based model suitable for use in a genetic screen to identify cellular factors that control prion-like propagation of tau aggregation.

### Genetic screen to identify cellular factors that control prion-like propagation of tau aggregation

In order to identify cellular factors that control propagation of tau aggregation (**Fig. 1A**), we conducted a CRISPR-interference (CRISPRi)-based genetic screen (**Fig. 2A**). First, we transduced the reporter cell line described above with a lentiviral expression construct for a catalytically inactive Cas9-BFP-KRAB (dCas9-BFP-KRAB) fusion protein. dCas9-BFP-KRAB can be directed by small guide RNAs (sgRNAs) to silence a gene of interest (22), enabling massively parallel genetic screens in mammalian cells (18). We then transduced the cells with pooled sgRNA libraries that target protein homeostasis factors, which we designed specifically for this study based on the rationale that protein homeostasis factors were likely to control or modulate tau aggregation and clearance. These libraries target 2,949 genes encoding genes that function in autophagy, protein folding, or the ubiquitin-proteasome system with at least five independent sgRNAs for each gene, plus 750 non-targeting control sgRNAs. Cells transduced with these libraries were exposed to recombinant tau fibrils at concentrations that would yield FRET positive cells at 50% of the maximum percentage of FRET positive cells (**Fig. 1I**), thereby maximizing the dynamic range for detecting cellular factors that either increase or decrease tau aggregation. FRET negative and FRET positive cell populations were separated by FACS, collecting sufficient numbers of cells from each population for an average 1000x representation (cells per sgRNA elements in the library). Genomic DNA was isolated and the locus encoding the sgRNAs was PCR-amplified. Frequencies for each sgRNA in each population were determined by next generation sequencing. We evaluated genes for the effect their knock-down had on the formation of tau aggregates (**Table S1**) using our previously described bioinformatics pipeline (18, 23-25).

**Figure 2.**
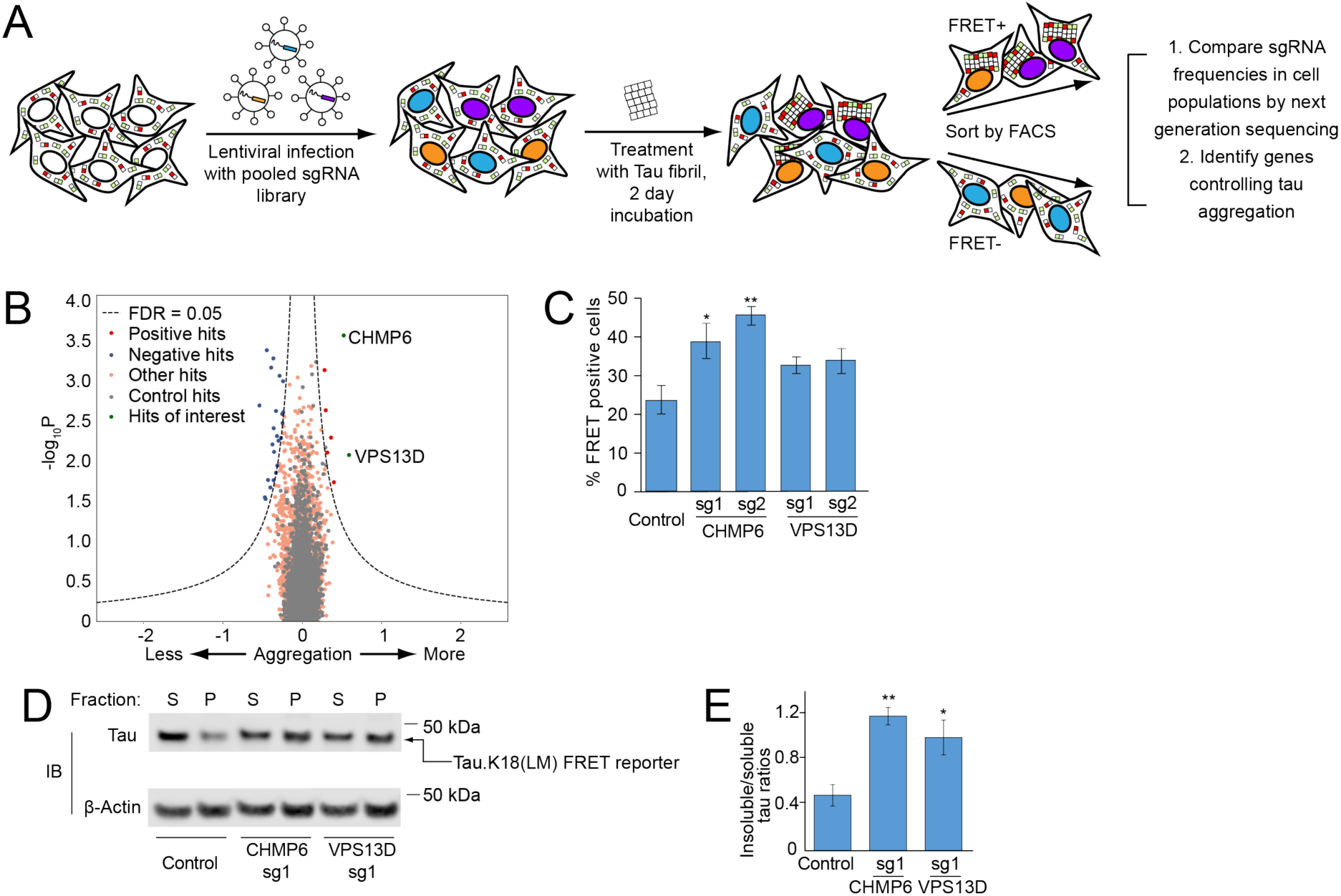
CRISPRi screen for cellular factors controlling tau aggregation. (**A**) Strategy for pooled FRET-based CRISPRi screen. FRET reporter cells stably expressing the CRISPRi machinery (dCas9-BFP-KRAB) were transduced with pooled lentiviral expression libraries of sgRNAs targeting proteostasis genes. Following transduction and selection, cells were treated with tau fibrils and incubated for 2 days. Cells were detached and sorted into FRET negative and positive populations by Fluorescence-Activated Cell Sorting (FACS). sgRNA-encoding cassettes were amplified from genomic DNA of the cell populations and their frequencies were quantified using next generation sequencing to identify genes that control tau aggregation. (**B**) Volcano plot summarizing phenotypes and statistical significance (by our MAGeCK-iNC pipeline, see Materials and Methods) of the genes targeted by the sgRNA libraries. Non-targeting sgRNAs were randomly grouped into negative control “quasi-genes” (grey dots) to derive an empirical false-discovery rate (FDR). Hit genes that passed an FDR < 0.05 threshold are shown in blue (knockdown decreases aggregation) or red (knockdown increases aggregation), other genes are shown in orange. Two hit genes of interest are shown in green and labeled. (**C-E**) Validation of hit genes CHMP6 and VPS13D. FRET reporter cells transduced with individual sgRNAs targeting two hit genes or a non-targeting control sgRNA, and 5 days after transduction treated for 2 days with tau fibrils. (C) % of FRET positive cells was quantified by flow cytometry. Error bars represent standard deviation of n=3 technical replicates. *P<0.05, **P<0.01 (two-tailed Student’s t test for comparison to the non-targeting control sgRNA). (D) Representative immunoblot for the tau-fluorescent protein construct in the soluble and insoluble fractions as in Fig. 1H. (E) Quantification of insoluble/soluble tau ratios from immunoblots in Fig 2D. Error bars represent standard deviation for n=3 biological replicates. P<0.05, **P<0.01, (two-tailed Student’s t test for comparison to the non-targeting control sgRNA).

Two genes stood out for the strong enhancement of tau aggregation by their knock-down: CHMP6 and VPS13D (**Fig. 2B**). We decided to prioritize these two genes for further characterization, since both are related to genes implicated in neurodegenerative diseases.

CHMP6 is part of the Endosomal Sorting Complex Required for Transport (ESCRT)-III complex, which is required for numerous cellular processes involving membrane remodeling (26). Mutations in the ESCRT-III component CHMP2B cause familial frontotemporal lobar dementia (FTD) and have been shown to cause endolysosomal defects (27, 28).

The *VPS13* protein family is comprised of four closely-related proteins, *VPS13A-D* (29). VPS13 family proteins are localized at various inter-organelle membrane contact sites and facilitates non-vesicular lipid transport (30, 31). Interestingly, mutations *VPS13D* are associated with recessive ataxia (32). Previously, *VPS13A* and *VPS13C* mutations have been associated with a Huntington’s-like syndrome (Chorea-Acanthocytosis) (33, 34) and Parkinson’s disease (35), respectively.

To confirm these screening hits, we cloned 2 individual sgRNAs each targeting CHMP6 and VPS13D, and confirmed target knockdown by qPCR (**Table S2**). Using these sgRNAs, we validated the effect of CHMP6 and, to a lesser extent, VPS13D knockdown on tau aggregation by flow cytometry (**Fig. 2C**) and biochemical solubility assay (**Fig. 2D, E**).

### CHMP6 knockdown accelerates tau aggregation following tau seed uptake

We next investigated the mechanism by which knockdown of CHMP6 and VPS13D might affect tau aggregation. First, we excluded the possibility that knockdown of these genes alters the levels of our tau reporter (**Fig. 3A**). Since we previously identified factors controlling cellular uptake of tau (16), we tested whether knockdown of CHMP6 and VPS13D impacted the uptake of tau fibrils. However, we found that knockdown of the genes did not impact tau fibril uptake (**Fig. 3B**), suggesting that their impact on tau aggregation is mediated downstream of seed uptake.

**Figure 3.**
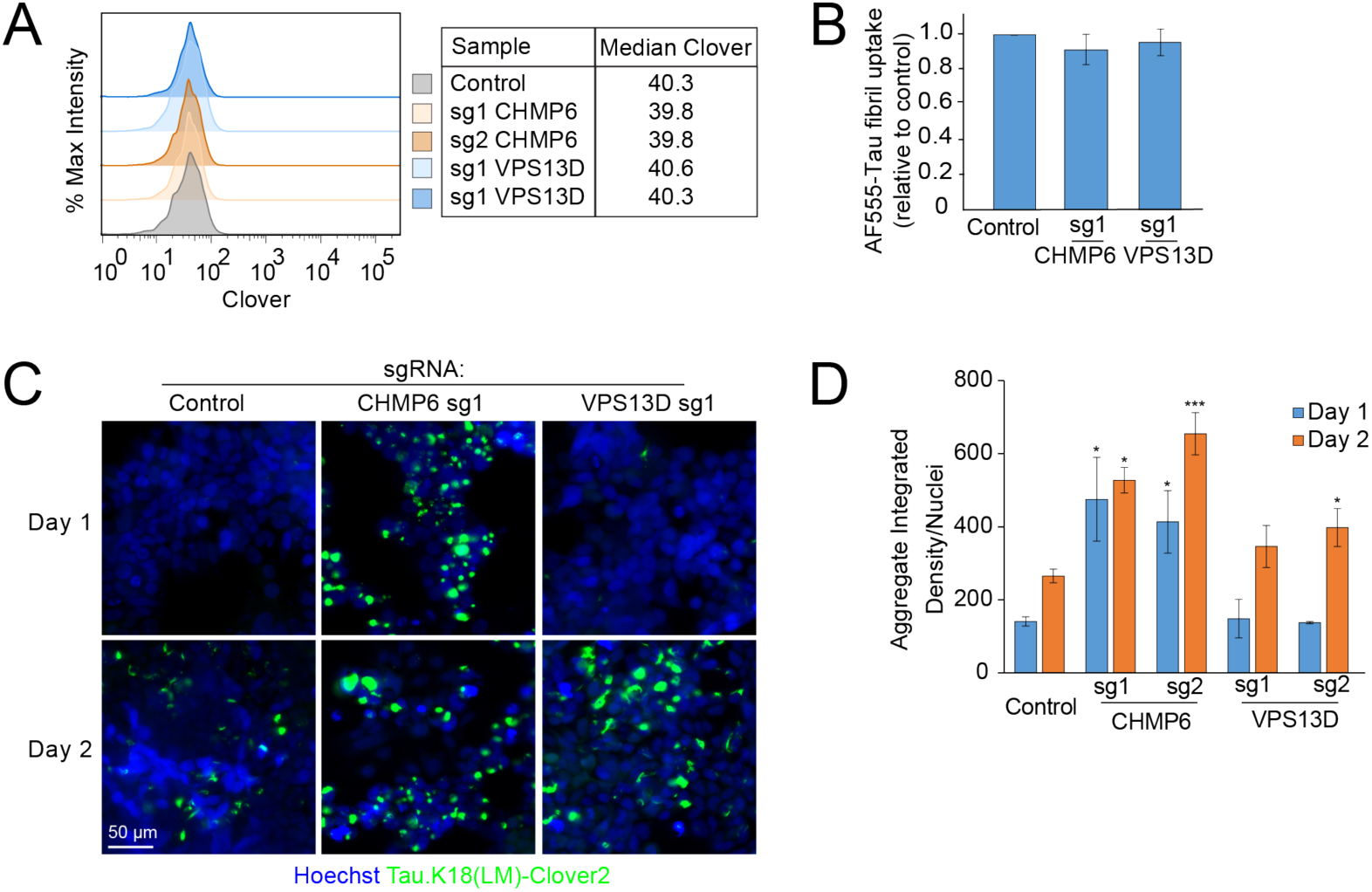
CHMP6 knockdown accelerates the prion-like propagation of tau aggregation. FRET reporter cells or CRISPRi-HEK293T cells were transduced with individual sgRNAs targeting CHMP6 or VPS13D, or a non-targeting control sgRNA, and characterized for different phenotypes 5 days after transduction (**A**) Knockdown of CHMP6 and VPS13D does not impact steady state levels of the tau-Clover2 construct in FRET reporter cells, as quantified by flow cytometry. (**B**) Individual gene knockdown does not impact uptake of tau fibrils. CRISPRi HEK293T cells were incubated with AF555-labeled tau fibrils for 1 hour at 37° C, stringently washed and red fluorescence representing internalized tau fibrils was quantified by flow cytometry. Bar graph shows normalized fluorescence intensities and standard deviation of n=3 technical replicates. (**C**) CHMP6 knockdown accelerates prion-like propagation of tau aggregation. Representative fluorescence micrographs of the Tau.K18(LM)Clover2 reporter in cells 1 and 2 days after fibril addition. Nuclei were counter-stained with Hoechst 33342. (**D**) Quantification of **Fig. 3C**. Tau aggregates were quantified by integrated density across the entire image and divided by total nuclei per image. Error bars represent standard deviation where n=3 images per condition (with at least 50 nuclei per image). *P<0.05, ***P<0.001 (two-tailed Student’s t test for comparison to the values for non-targeting control sgRNA of the same day).

We next sought to evaluate whether knockdown of CHMP6 and VPS13D increased the rate of tau aggregation, or decreased the rate of tau aggregate clearance. To this end, we utilized high-content imaging analysis to track the fibril-induced aggregation of tau over time. As expected from the results in our primary screen, we observed increased levels of aggregates 48 hours post seeding with fibrils when either CHMP6 or VPS13D were knocked down (**Fig. 3C,D**). Intriguingly, the timeline of tau aggregate formation was differentially affected by the two gene knockdowns. While VPS13D knock-down did not cause a statistically significant increase in aggregates 24 hours after treatment with tau fibrils compared to a non-targeting control sgRNA, CHMP6 knockdown promoted early aggregation 24 hours post seeding (**Fig. 3C,D**). Interestingly, aggregate formation from 24 to 48 hours post-seeding did not change substantially with CHMP6 knockdown, suggesting that the majority of soluble tau rapidly aggregates following treatment with tau fibrils in that background. While we observe a rapid increase in aggregate formation in CHMP6 knockdown cells, this does not rule out an additional effect of CHMP6 knockdown on aggregate clearance. Given the intriguing acceleration of tau aggregation by CHMP6, and the comparatively weaker phenotype of VPS13D, we decided to focus our mechanistic studies on CHMP6.

### Endolysosomal escape of tau seeds is rate limiting for propagation of tau aggregation

To investigate the mechanism by which CHMP6 knockdown accelerates seeded tau aggregation, we monitored fibril entry and aggregate formation simultaneously by longitudinal imaging in cells expressing a non-targeting control sgRNA compared to CHMP6 knockdown. (**Fig. 4A, Supplementary Movie 1** and **2**). In CHMP6 knockdown cells, large tau aggregates rapidly formed soon after tau fibrils localized to cells, within 12 hours post seeding. In control cells, by contrast, tau fibrils localized to control cells long before aggregates form. We confirmed that these fibril puncta partially colocalize with the late-endosome/lysosome markers LAMP1 and Rab7a, and frequently localize to the lumen of LAMP1 and Rab7a positive compartments (**Fig. 4B, Fig. S1A, Supplementary Movies 3,4)**. (Large LAMP1 and Rab7a vesicles with visible lumina were observed also in the absence of fibrils, and therefore not induced by the fibrils themselves, Fig. S1B). These results suggest that fibrils normally accumulate in endolysosomal compartments, where they do not encounter cytosolic tau. In CHMP6 knockdown cells, colocalization of fibrils with LAMP1-positive compartments was markedly reduced (**Fig. 4B,C and Supplementary Movie 5**). Therefore, CHMP6 knockdown seems to accelerate exit of fibrils from the endolysosomal pathway into the cytosol, where it can then seed aggregation of cytosolic tau.

**Figure 4.**
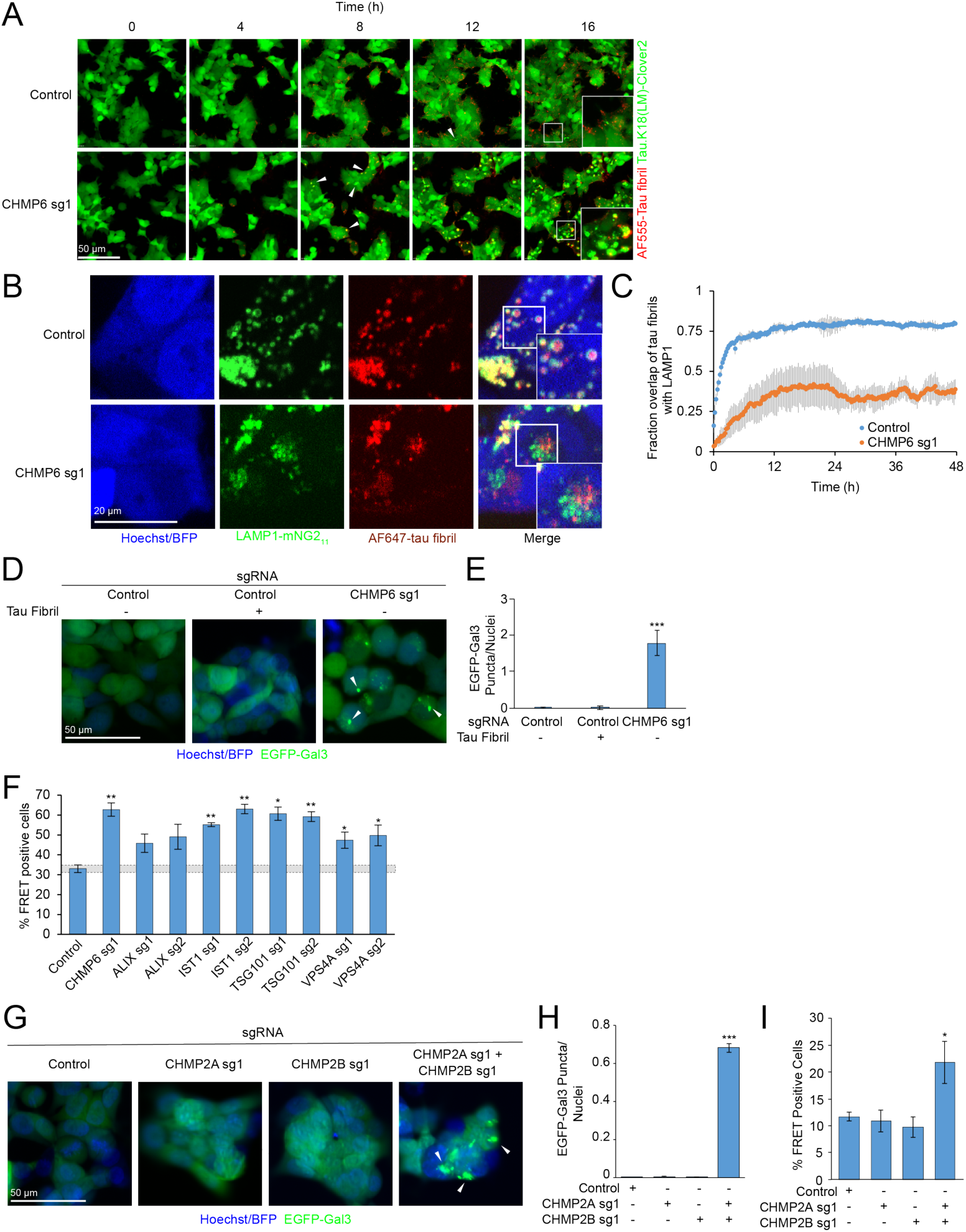
CHMP6 knockdown compromises endolysosomal membrane integrity. (**A**) Time-lapse microscopy of cell entry of tau-AF555 fibrils and resulting aggregation of the cytosolic tau-Clover2 construct. Representative images of CRISPRi-HEK293T cells expressing tau.K18(LM)-clover2 transduced with either non-targeting control (top) or *CHMP6*-targeting sgRNA (bottom) are shown. Corresponding movies are provided as Supplementary Movies 1 (control sgRNA) and 2 (CHMP6 sgRNA). (**B**) Representative fluorescence microscopy images of CRISPRi-HEK293T cells LAMP1-mNG_11_ endogenously labeled with the split-mNeonGreen system. Cells were transduced with non-targeting control (top) or CHMP6 sgRNA (bottom) and treated with AF647-tau fibrils and followed by automated time-lapse microscopy for 48 hour time interval. Images for the 12 h timepoint are shown here. Corresponding movies for 48 hour time intervals are provided as Supplementary Movies 4 (Control sgRNA) and 5 (CHMP6 sg1) (**C**) Quantification of Tau fibril colocalization with LAMP1 from representative images and movies shown in figure Fig 4B and Supplementary Movies 4 and 5. Time represents start of treatment with tau fibrils and image acquisition. Error bars represent standard deviation for n=3 technical replicates (with at least 5 nuclei per image). (**D**) CHMP6 knockdown causes endolysosomal vesicle damage. Representative fluorescence microscopy images of CRISPRi-HEK293T cells expressing an EGFP-Galectin3 (EGFP-Gal3) reporter transduced with either control (*left*) or CHMP6 (*right*) sgRNA. Nuclei were counter-stained with Hoechst 33342. (**E**) Quantification of EGFP-Gal3 puncta divided by number of nuclei in fluorescence microscopy images shown in **Fig 4D**. Error bars represent standard deviation for n=3 technical replicates (with at least 50 nuclei per image). ***P<0.001 (two-tailed Student’s t test for comparison to the non-targeting control sgRNA). (**F**) Knockdown of various ESCRT components increases tau aggregation. FRET reporter cells were transduced with individual sgRNAs targeting ESCRT components or a non-targeting control sgRNA, and 5 days after transduction treated for 2 days with tau fibrils. Error bars represent standard deviation for n=3 technical replicates. *P<0.05, **<0.01 (two-tailed Student’s t test for comparison to the non-targeting control sgRNA). (**G**) Simultaneous, but not individual, knockdown of CHMP2A and CHMP2B results in endolysosomal damage, monitored as in Fig. 4C. Nuclei were counter-stained with Hoechst 33342. Scale bar = 50 µm (**H**) Quantification of EGFP-Gal3 puncta divided by nuclei in fluorescence microscopy images shown in Fig. 4G. Error bars represent standard deviation for n=3 technical replicates (with at least 50 nuclei per image). ***P<0.001 (two-tailed Student’s t test for comparison to the non-targeting control sgRNA). (**I**) Simultaneous, but not individual, knockdown of CHMP2A and CHMP2B increases prion-like tau aggregation. % FRET positive reporter cells transduced with sgRNAs as indicated 2 days after tau fibril treatment. Error bars represent standard deviation where n=3 technical replicates. *P<0.05 (two-tailed Student’s t test for comparison to the non-targeting control sgRNA).

A mechanism underlying this CHMP6 phenotype is suggested by the recently reported role of the ESCRT machinery in the repair of endolysosomal membrane damage (36). We hypothesized that knocking down CHMP6 may compromise ESCRT-mediated membrane repair and facilitate tau fibril escape from damaged endolysosomes. We tested this hypothesis by monitoring the formation endolysosomal damage using a cytosolic GFP fusion of galectin 3 (GAL3), a lectin that binds β-galactosides and forms puncta when these sugars are exposed on damaged endolysosomes (37). While tau fibrils themselves did not cause measurable endolysosomal damage based on our GAL3-GFP reporter (**Fig. 4D,E**), knocking down CHMP6 indeed induced GAL3-GFP puncta, revealing endolysosomal damage. This demonstrates that CHMP6 plays a critical role in the maintenance of endolysosomal integrity.

CHMP6 was the only ESCRT protein that was a strong hit in our primary CRISPRi screen. However, targeted knockdown of several other ESCRT components, including members of the ESCRT-I complex (Tsg101), the ESCRT-III complex (Ist1) and the ESCRT-III associated Vps4a promoted tau aggregation (**Fig. 4F**), pointing to a role for the ESCRT pathway in general, as opposed to a specialized role for CHMP6.

Other ESCRT proteins have several paralogues in the human genome, so we hypothesized that these paralogues may have been false-negatives in the CRISPRi screen because they have partially redundant functions. To test this hypothesis, we targeted CHMP2B, a gene with disease-associated mutations involved in familial frontotemporal lobar dementia (38). We hypothesized that its phenotype might have been masked by its close homolog, CHMP2A, which could partially compensate for a loss in CHMP2B function. Indeed, we found that simultaneous, but not individual, knockdown of CHMP2A and CHMP2B generated GAL3 puncta indicative of endolysosomal damage (**Fig. 4G,H**) and likewise promoted prion-like propagation of tau aggregation (**Fig. 4I**). This finding supports our hypothesis that maintenance of endolysosomal membrane integrity by the broader ESCRT-machinery counteracts endolysosomal escape of tau seeds.

A key implication of this model is that endolysosomal damage may promote the prion-like propagation of tau aggregation. To test this concept, we treated cells with Leucyl-Leucyl-o-Methyl-ester (LLOME), a lysosomotropic compound that accumulates in acidified organelles and rapidly forms membranolytic polymers after cleavage by cathepsin C (36, 39, 40). We confirmed that LLOME damages endolysosomal membranes in our cell line using the GAL3-GFP reporter (**Fig. 5A,B**). Interestingly, LLOME phenocopied CHMP6 knockdown in its acceleration of seeded tau aggregation only at concentrations where we observe endolysosomal membrane damage (**Fig. 5C**).

**Figure 5.**
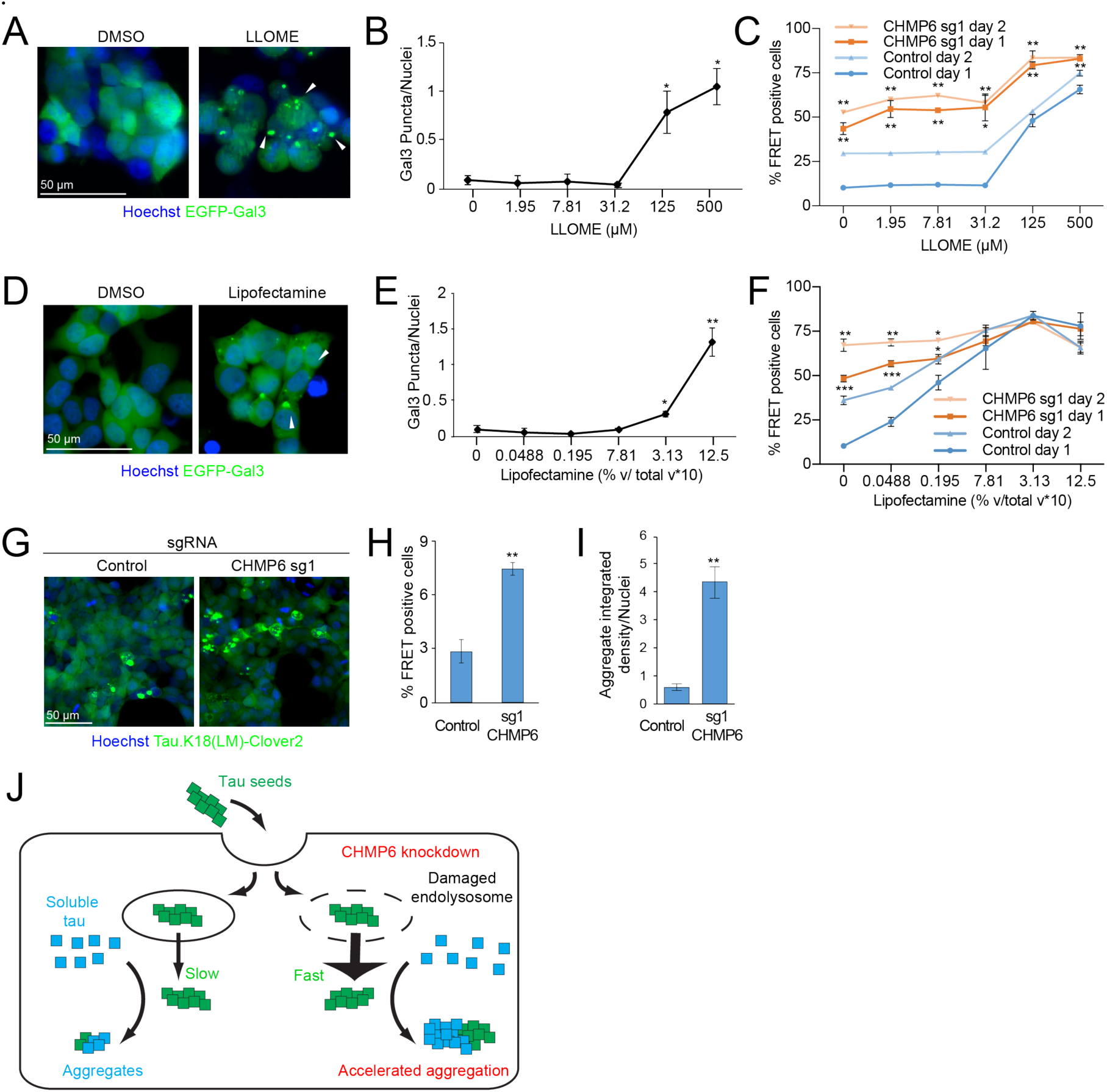
Small molecules damage endolysosomal compartments and phenocopy the acceleration of the prion-like propagation of tau aggregation following CHMP6 knockdown. (**A**) Treatment with the lysosomotropic drug leucyl-leucyl-O-methyl-ester (LLOME) damages endolysosomal vesicles. Representative fluorescence microscopy images of CRISPRi-HEK293T cells expressing the EGFP-Gal3 reporter treated with DMSO (left) or 500 µM LLOME (right) for 24 h. (**B**) Quantification of EGFP-Gal3 puncta divided by number of nuclei in fluorescence microscopy images shown in **Fig 5A**. Error bars represent standard deviation for n=3 technical replicates (with at least 50 nuclei per image). ***P<0.001 (two-tailed Student’s t test for comparison to DMSO control). (**C**) LLOME treatment accelerates the prion-like propagation of tau aggregation. % FRET positive cells transduced with control (*blue*) or CHMP6 sgRNA (*orange*) were analyzed 24 (*dark line*) or 48 (*light line)* hours following co-treatment with DMSO or increasing concentrations of LLOME and tau fibrils. Error bars represent standard deviation for n=3 technical replicates. *P<0.05, **P<0.01 (two-tailed Student’s t test for comparison to the values for non-targeting control sgRNA of the same day) (**D**) Lipofectamine treatment damages endolysosomal vesicles. Representative fluorescence microscopy images of CRISPRi-HEK293T cells expressing the EGFP-Gal3 reporter treated with DMSO (left) or 1.25 % v/v Lipofectamine 2000 (right) for 6 h. (**E**) Quantification of EGFP-Gal3 puncta divided by number of nuclei in fluorescence microscopy images shown in **Fig 5D**. Error bars represent standard deviation for n=3 technical replicates. ***P<0.001 (two-tailed Student’s t test for comparison to no lipofectamine treatment). (**F**) Lipofectamine treatment accelerates the prion-like propagation of tau aggregation. % FRET positive cells transduced with control (*blue*) or CHMP6 sgRNA (*orange*) were analyzed 24 (*dark line*) or 48 (*light line)* hours following co-treatment with PBS or increasing concentrations of lipofectamine 2000 and tau fibrils. Error bars represent standard deviation where n=3 technical replicates. *P<0.05, **P<0.01, ***P<0.001 (two-tailed Student’s t test for comparison to the values for non-targeting control sgRNA of the same day) (**G-I**) CHMP6 knockdown increases the prion-like propagation tau aggregation when seeding with Alzheimer’s patient brain extracts. (G) Representative images of FRET reporter cells transduced with control (*left*) or CHMP6 sgRNA (*right*) and treated with Alzheimer’s patient brain extract after 5 days (H) Quantification of images in **Fig. 5G**. Tau aggregates were quantified by integrated density across the entire image and divided by total nuclei per image. Error bars represent standard deviation for n=3 images per condition (with at least 50 nuclei per image). **P<0.01 (two-tailed Student’s t test for comparison to non-targeting control sgRNA) (I) % FRET positive cells 5 days following treatment with Alzheimer’s patient brain extract. Error bars represent standard deviation where n=3 technical replicates. **P<0.01 (two-tailed Student’s t test for comparison to non-targeting control sgRNA). (**J**) Model for the increased tau aggregation via CHMP6 knockdown and generation of damaged endolysosomes.

As mentioned above, lipocationic reagents, such as Lipofectamine, are frequently used to deliver tau aggregates to cells for in vitro studies of prion-like propagation (13, 14, 41). Interestingly, these agents have previously been demonstrated to induce endolysosomal damage (42). Thus, lipocationic agents might facilitate cargo delivery and escape, in part, by causing endolysosomal membrane damage. Indeed, we found that GAL3 puncta indicative of endolysosomal damage are visible 24 hours after treating cells with Lipofectamine (**Fig. 5D,E)**. Moreover, pre-treatment with lipofectamine 6 hours prior to seeding cells with tau fibrils significantly increased the formation of tau aggregates in a concentration-dependent manner, including at concentrations lower than the threshold required to induce a Gal3 reporter response (**Fig. 5F**). This suggests that lipofectamine may assist in the prion-like spread of tau aggregates both by acting as a delivery vehicle and damaging endolysosomal membranes.

When we combined CHMP6 knock-down with either LLOME or lipofectamine treatment, we found that the relative impact of CHMP6 knockdown on tau aggregate seeding was diminished in the presence of LLOME (**Fig. 5C**) or lipofectamine (**Fig. 5F**), supporting the idea that CHMP6 knockdown and LLOME/lipofectamine treatment promote propagation of tau aggregation at least partially via the same mechanism.

Finally, we wanted to validate the CHMP6 phenotype using tau seeds derived from Alzheimer’s patient brain-derived extracts. Indeed, we found that CHMP6 knockdown increased the rate of tau aggregation in our reporter line seeded with patient brain-derived tau by both microscopy (**Fig. 5G,H**) and flow cytometry (**Fig. 5I**). Taken together, our results support endolysosomal escape of tau seeds as a rate-limiting step in our cell-based model of prion-like tau propagation. Propagation can be accelerated by compromising endolysosomal integrity either by directly damaging endolysosomes or by interfering with their repair through the ESCRT machinery (**Fig. 5J**).

## DISCUSSION

Using our CRISPRi-based genetic screening platform in a cell-based model of prion-like propagation of tau aggregation, we found that defects in the ESCRT machinery compromise the integrity of the endolysosomal pathway and thereby promote endolysosomal escape of tau seeds and accelerated propagation of tau aggregation. While our observations were made in a cell-based model, it is intriguing to speculate that they are relevant for propagation of tau aggregation in the context of neurodegenerative diseases. Indeed, endolysosomal changes are among the first cellular symptoms in Alzheimer’s Disease (43), and have been postulated to be a central driver of pathogenesis in many neurodegenerative diseases (44-46). Furthermore, several risk genes for neurodegenerative diseases are thought to function in the endolysosomal pathway, including CHMP2B (38).

While the ESCRT-III subunit CHMP6 was a top hit in our genetic screen, CHMP2B knockdown by itself did not have a major impact on endolysosomal integrity and the propagation of tau aggregation. This was likely the case because human cells express CHMP2A, a paralogue of CHMP2B which can mostly compensate for CHMP2B in our cell-based model – combined knockdown of CHMP2A and CHMP2B phenocopied CHMP6 knockdown. CHMP6 does not have a paralogue in mammalian cells and is an essential gene, whereas CHMP2B is non-essential, based on the Cancer dependency map, depmap.org (47), and knock-out mouse phenotypes (48). This provides a rationale for CHMP6 deficiency not being associated with neurodegenerative diseases – it may not be compatible with life. CHMP2B deficiency can be expected to cause a milder phenotype that is unmasked only later in life.

Many mechanisms have been proposed to explain the toxicity of tau aggregates. Intriguingly, tau aggregates can damage membranes in vitro (49), and may damage the endolysomal pathway in patient neurons. In combination with our findings, such a mechanism of toxicity would predict a “vicious circle” or feed-forward mechanism, in which tau aggregates would damage the endolysosomal pathway, thereby promoting their own propagation. They could also promote spreading of other aggregates, compatible with the co-occurrence of different protein pathologies, such as tau and a-synuclein, in many cases of neurodegenerative disease (50). However, in our HEK293T cell-based model, we did not find evidence of endolysosomal damage introduced by tau fibrils (Fig. 4D,E) – possibly due to differential susceptibility of different cell types to tau toxicity.

In summary, our results further support the concept that therapeutic strategies aimed at maintaining or restoring the function of the endolysosomal pathway or promoting its repair may be promising in neurodegenerative diseases. Future studies will be aimed at understanding mechanisms underlying the VPS13D phenotype, which seems distinct from the endolysosomal escape pathway controlled by CHMP6.

While our genetic screen with libraries targeting protein homeostasis factors unexpectedly uncovered the ESCRT machinery in counter-acting endolysosomal escape of tau, we had expected to find molecular chaperones or cochaperones controlling tau aggregation among the top hits. Results obtained *in vitro* (51-54) and *in vivo* (55) suggest that specific chaperones and co-chaperones can strongly modulate tau aggregation, and are potential therapeutic targets for tauopathies (56). The fact that knockdown of individual chaperones did not have a major impact on tau aggregation could be due to redundancy in the chaperone network of cells. Future CRISPR activation (CRISPRa) screens have the potential to yield complementary results by over-expressing endogenous genes (18, 19) and may reveal relevant chaperones in the cellular context.

Finally, future screens in iPSC-derived neurons using our recently developed platform (25) may reveal additional, neuron-specific pathways, and also uncover factors that underlie selective vulnerability of specific neuronal sub-types (57).

## EXPERIMENTAL PROCEDURES

### Preparation of extracts from Alzheimer’s Disease patient brains

The Alzheimer’s Disease brain sample was received from the Neurodegenerative Brain Bank of the UCSF Memory and Aging Center (UCSF/MAC). All research participants at UCSF/MAC undergo longitudinal clinical and imaging assessment. Upon death, the fresh brain was slabbed into 8- to 10-mm thick coronal slabs upon procurement. These slabs were alternately fixed, in 10% neutral buffered formalin for 72 hours, or snap frozen. Twenty-six tissue blocks covering dementia-related regions of interest were dissected from the fixed slabs, and hematoxylin and eosin and immunohistochemical stains were applied following standard diagnostic procedures developed for patients with dementia (58, 59). Immunohistochemistry was performed using antibodies against TDP-43 (rabbit, 1:2000, Proteintech Group, Chicago, IL, USA), hyperphosphorylated tau (CP-13, S202/T205, mouse, 1:250, courtesy of P. Davies), beta-amyloid (1-16, mouse, clone DE2, 1:500, Millipore, Billerica, MA, USA), alphasynuclein (LB509, mouse, 1:5000, courtesy of J. Trojanowski and V. Lee). All immunohistochemical runs included positive control sections to exclude technical factors as a cause of absent immunoreactivity. Neuropathological diagnosis followed currently accepted guidelines (60-64). For this study, a region from the parietal cortex containing high amount of AD-tau pathology was sampled from a snap frozen block. A brain extract was prepared and phosphotungstate-insoluble fractions were purified as previously described (13). The extract was diluted in PBS 1:40 in DPBS and flash frozen in liquid nitrogen and stored at −80 °C.

### Purification, characterization, labeling, and fibrillization of recombinant tau

Human WT 0N4R-6xHis tau protein was expressed in Rosetta™ 2(DE3) competent cells (MilliporeSigma #71400-3) essentially as previously described (51). Briefly, protein expression was induced with 200 µM IPTG for 3.5 hours at 30 °C. Cells were lysed via a microfluidizer (Microfluidics Cat# M-100EH) followed by boiling of the lysate for 20 min. The clarified supernatant was subsequently dialyzed overnight into Buffer A (20 mM MES pH 6.8, 50 mM NaCl, 1mM EGTA, 1 mM MgCl2, 2 mM DTT, 0.1 mM PMSF) and purified by cation exchange using a HiTrap Capto SP ImpRes column (GE Cat# 17546851) with elution buffer (Buffer A with 1 M NaCl). Fractions containing tau as determined by Coomassie-stained SDS-PAGE were dialyzed into PBS, concentrated with an Amicon Ultra-15 centrifugal 3 kDa MWCO filter (Millipore Cat# UFC900324), endotoxin purified using a Pierce high capacity endotoxin removal spin column (ThermoFisher Cat# 88274), filter sterilized using a Millex-GV syringe filter unit (Millipore Cat# SLGV033RB), and snap frozen in PBS at - 80 °C. Aggregation was induced by incubating 10 µM tau (0.43 mg/mL) with .022 mg/mL heparin (Fisher Cat# 9041-08-1, lot# 177772) and shaken at 37 °C overnight in a shaker at 1200 rpm (VWR Cat#12620-942).

To generate fluorescently labeled tau fibrils, 0.6 µl 10 mg/mL Alexa Fluor-555 (ThermoFisher Cat# A37571), 180 µl of 0.43 mg/mL tau fibrils, and 19.4 µl 1M sodium bicarbonate were mixed at room temperature in the dark for 1h. Labeled tau fibrils were subsequently purified from unlabeled dye with using a Zeba 7k MWCO spin desalting column (ThermoFisher Cat# 89882).

Tau fibrils were negatively stained with 0.75% uranyl formate (pH 5.5-6.0) on thin amorphous carbon layered 400-mesh copper grids (Pelco Cat# 1GC400). Five µL of sample was applied to the grid for 20s before taking the droplet off with Whatman paper, followed by two washes with 5 µL ddH2O and three applications of 5 µL uranyl formate removed by vacuum. Grids were imaged at room temperature using a Fei Tecnai 12 microscope operating at 120kV. Images were acquired on a US 4000 CCD camera at 66873x resulting in a sampling of 2.21 Angstrom/pixel.

### Plasmid and library design and construction

Plasmids for the FRET-based aggregation reporter were constructed by cloning a fusion of the K18 repeat domain of tau containing the P301L/V337M mutation (20) in frame with C-terminal Clover2 (Addgene #54711) or mRuby2 (Addgene #54768, (21), gifts from Michael Davidson, into the lentiviral expression vector pMK1200 (23), Addgene #84219) under the control of the constitutive EF1A promoter, to obtain pMK1253 or pMK1254, respectively. The K18-Scarlet-I reporter was constructed by cloning a fusion of the K18 construct from pMK1253 in frame with the C-terminal mScarlet-I (Addgene #98839, a gift from Dorus Gadella (65)) as well as replacing the EF1a promoter with a CAG promoter to obtain pJC49.

Pooled CRISPRi sgRNA libraries targeting human protein homeostasis genes were designed using our next-generation algorithm (66). SgRNA protospacers for these libraries are listed in Table S2. Oligonucleotide pools encoding the library were synthesized by Agilent, PCR amplified and cloned into our optimized lentiviral sgRNA expression vector as previously described (18).

For generation of individual sgRNAs, pairs of oligonucleotides (IDT) were annealed and ligated into our optimized lentiviral sgRNA expression vector. For double sgRNA expression constructs, CHMP2B and CHMP2A targeting oligos were annealed and ligated into pMJ114 and pMJ179, and a double-sgRNA vector was generated from these as previously described (67).

The fluorescent Gal3 reporter was PCR amplified from pEGFP-hGal3 (Addgene #73080), a gift from Tamotsu Yoshimori) and Gibson cloned into pJC41, which uses the pMK1200 backbone (described above) and replaces the EF1a promoter with a CAG promoter.

For stable expression of the CRISPRi machinery, we modified our established lentiviral (d)Cas9 expression vectors (18) by replacing the SFFV promoter with a minimal ubiquitous chromatin opening element (UCOE) (68) upstream of the EF1alpha promoter, resulting in pMH0006 (UCOE-SFFV-dCas9-BFP-KRAB).

### Cell culture, cell line generation, and treatment conditions

All cells were maintained in a tissue culture incubator (37 °C, 5% CO_2_) and checked regularly for mycoplasma contamination. HEK293T cells were cultured in DMEM supplemented with 10% fetal bovine serum (Seradigm Cat# 97068-085, Lot# 076B16), Pen/Strep (Life Cat# 15140122), and L-glutamine (Life Cat# 25030081).

To generate the FRET reporter line, HEK293T cells were infected with lentivirus from plasmids pMK1253 and pMK1254 and cells with the highest dynamic FRET signal 2 days after seeding with tau fibrils were selected. To introduce CRISPRi functionality, the cells were lentivirally transduced with pHR-SFFV-dCas9-BFP-KRAB (Addgene #46911, a gift from Stanley Qi and Jonathan Weissman (22)), mono-clonal cell lines were selected and CRISPRi activity was validated as previously described (25).

Cellular markers were endogenously labeled using the split-mNeonGreen2 system (69), following conditions described in (70). Briefly, synthetic guide RNAs (IDT, Alt-R reagents) were first complexed in vitro with purified *S. Pyogenes* Cas9 protein (UC Berkeley Macrolab). Cas9/RNA complexes were then mixed with single-stranded DNA oligo donors (IDT, Ultramer reagents) and nucleofected (Lonza Cat#AAF-1002B, Amaxa program CM-130) into HEK cells stably expressing SFFV-mNeon-Green2_1-10_. Fluorescent cells were selected by flow cytometry (SONY biotechnology Cat# SH800S). Sequences for CRISPR RNA and donors used are listed as follows: LAMP1 (C-term mNG11): crRNA sequence 5’-GTGCAC-CAGGCTAGATAGTC-3’; donor oligonucleotide sequence 5’-CCCAGA-GAAAGGAACAGAGGCCCCTGCAGCTGCTGTGCCTGCGTGCACCAGGCTACATCATATCGGTAAAGGCCTTTTGCCACTCCTTGAAGTTGAGCTCGGTACCACTTCCTGGACCTTGAAACAAAACTTCCAATCCGCCACCGATAGTCTGGTAGCCTGCGTGACTCCTCTTCCTGCCGACGAGGTAGGCGATGAGG-3’; RAB7A (N-term mNG11): crRNA sequence 5’-TAGTTTGAAGGATGACCTCT-3’; donor oligonucleotide sequence 5’-TGTTTCCATCACACTCACAGTGATTTCTCCTTTTCCCCCTTTAGTTTGAAGGATGACCGAGCTCAACTTCAAGGAGTGGCAAAAGGCCTTTACCGATATGATGGGTGGCGGATTGGAAGTTTTGTTTCAAGGTCCAGGAAGTGGTACCTCTAGGAAGAAAGTGTTGCTGAAGGTTATCATCCTGGGAGATTCTGGGTAAG-3’.

To generate the LAMP1/K18-mScarletI CRISPR-cells, the LAMP1-mNG_11_ cells were lentivirally transduced and sorted sequentially with pMH006 and pJC49.

To generate CRISPRi-HEK293T cells that monitor EGFP-Gal3 damage or only generate tau.K18(LM)-Clover2 aggregates, CRISPRi-HEK293T cells were lentivirally transduced with pMK1253 and pJC41, and a polyclonal population was sorted by FACS.

### Primary CRISPRi screen

For pooled screening of libraries, 7.5 million HEK293T cells were seeded into a 15 cm^2^ plate with complete DMEM on day 0. On day 1, 5 µg of lentiviral plasmid packaging mix (24) and 5 µg of pooled sgRNA library plasmid was transfected using lipofectamine 2000 (ThermoFisher Cat# 11668019) and incubated for 2 days. On day 3, conditioned media was removed and filter sterilized using a Millex-GV syringe filter unit (Millipore Cat# SLGV033RB). Lentivirus was precipitated (Alstem Cat# VC100) according to manufacturer protocols and resuspended in complete DMEM. 20 million FRET reporter cells were added to lentivirus-containing media and seeded into a T175 flask. On day 4, media from the T175 was replaced with DMEM complete with 2.5 µg/mL puromycin. On day 8, Cells infected with pooled sgRNA libraries were trypsinized and replated into at 100 µl per well (25,000 cells/well) of several 96-well plate. In addition, 0.3 µl of 0.43 µg/µL of tau fibrils were added to each well. 48 hours later, cells were trypsinized and sorted using an Aria II FACS cytometer into FRET negative and FRET positive populations. Genomic DNA was isolated using a Macherey-Nagel Blood L kit (Machery-Nagel Cat# 740954.20) and followed according to manufacturer protocols. SgRNA-encoding regions were then amplified and sequenced as previously published (18). Phenotypes and P values for each gene were calculated using our most recent bioinformatics pipeline (https://kampmannlab.ucsf.edu/mageck-inc (25)). For genes targeted by more than one sgRNA library, values for the more significant phenotype were selected. Full results are listed in Table S1.

### Secondary assays based on microscopy and flow cytometry

To monitor tau aggregation, FRET reporter cells were seeded (25,000 cells/well) into 100 µL per well in a 96 well black bottom plates (Greiner Bio-One #655097) with 0.3 µL 0.43 mg/mL Tau fibrils on day 1 and analyzed 24 or 48 hours after seeding. For Alzheimer’s patient brain extracts, 1.5 µL extract, 0.375% total v/v Lipofectamine 2000 (ThermoFisher Cat# 11668019), and 7.85 µL OptiMEM (Thermo Cat# 31985062) were mixed and incubated at room temperature for 2 hours. Lipofectamine-brain extract complexes were then added to cells previously plated in 100 µL (10,000 cells/well) for 6 hours. Cells were analyzed 5 days after seeding. For analysis, cells were stained with Hoechst 33342 (ThermoFisher Cat# 5553141) at 1 µg/mL and analyzed by flow cytometry using a BD FACSCelesta or by fluorescence microscopy using an InCell 6000 (GE Cat# 28-9938-51). Digital images were analyzed using CellProfiler by counting the integrated density of identified aggregates/nuclei and averaged between 3 images. Cells with sgRNA knockdown were similarly analyzed using a comparable protocol 5 days after transduction with individual sgRNA-encoding lentivirus.

For experiments measuring tau aggregation in the presence of inducers of endolysosmal damage, FRET reporter cells were seeded (25,000 cells/well) into 100 µL per well in a 96 well black bottom plate and treated with LLOME (Sigma Cat# L7393-500MG) at varying concentrations with 0.3 µL 0.43 µg/µL Tau fibrils. For treatment with lipofectamine 2000, FRET reporter cells were seeded (25,000 cells/well) into 100 µL per well in a 96 well black bottom plate and treated with Lipofectamine 2000 at varying concentrations. Cells were then treated 0.3 µL 0.43 µg/µL Tau fibrils 6 hours later. 24 or 48 hours later after seeding, cells were stained with Hoechst 33342 (1 µg/mL) and analyzed by flow cytometry using a BD FACSCelesta or by fluorescence microscopy using an InCell 6000 GE (Cat# 28-9938-51). Digital images were collected and analyzed using CellProfiler by quantifying the integrated density of identified aggregates and Hoechst-stained nuclei. In cases where CellProfiler was unable to identify nuclei, nuclei were counted interactively using ImageJ. Cells with sgRNA knockdown were similarly analyzed using a comparable protocol 5 days after transduction with individual sgRNA-encoding lentivirus.

To monitor tau fibril uptake, on day 0, CRISPRi-HEK293T cells previously transduced for 5 days with lentivirus expressing single sgRNAs were seeded (25,000 cells/well) into 100 µL per well in a 96-well plate. On day 1, cells were treated with 1 µL 0.39 µg/µL AF555-tau fibril for 1 hr at 37 °C and collected for analysis by flow cytometry. Median mRuby2 values were calculated in FlowJo and averaged between 3 technical replicates.

To monitor tau.K18(LM)-Clover2 steady-state levels, on day 0, FRET reporter cells previously transduced for 5 days with lentivirus expressing single sgRNAs were seeded (25,000 cells/well) into 100 µL per well in a 96-well plate. On day 1, cells collected for analysis by flow cytometry. Median Clover2 values were calculated in FlowJo and averaged between 3 technical replicates.

To monitor localization of AF555-labeled tau fibrils, HEK293T cells expressing Tau.K18(LM)-Clover2 were seeded (12,500 cells/well) into 100 µL per well in a 96-well black bottom plate (Greiner Bio-One #655097) on day 0 in complete DMEM. On day 1, 0.3 µL 0.39 µg/µL AF555-tau fibrils were added to cell culture media and placed into an InCell 6000 (GE Cat# 28-9938-51) incubator. Images were taken by at 20 minute intervals between incubations.

To monitor co-localization of tau fibrils with LAMP1, fluorescently labeled CRISPRi-HEK293T cells were seeded in glass-bottom 96-well plates (Cellvis #P96-1.5P) pre-coated with fibronectin (Roche Cat# 11051407001) at 15,000 cells cells/well in 150 µL complete DMEM media (including 10% FBS). After incubation for 3 hours to allow for cell adhesion, cells were treated with 0.11 µg of AF555-tau PFFs per well. 22 hours post-treatment, cells were counter-stained with Hoechst 33342 (0.5 µg/mL, 30 min at 37 °C) and imaged in complete DMEM without phenol-red. Live-cell imaging was performed on a Dragonfly spinning-disk instrument (Andor) at 37 °C in 5% CO2 atmosphere equipped with a 63x/1.47 NA objective (Leica) and an iXon Ultra 888 EMCCD camera (Andor), acquiring time-lapse datasets at 0.4Hz. Images were analyzed by thresholding LAMP1 and tau fibril images and masking pixels positive for LAMP1 and tau fibrils. Colocalization was calculated by dividing the total image intensity of the masked image with total image intensity of the thresholded tau fibrils. Cell profiler scripts are available at kampmannlab.ucsf.edu/resources.

To monitor Gal3-EGFP puncta formation, CRISPRi-HEK293T cells expressing EGFP-Gal3 were seeded into 100 µL per well (25,000 cells/well) in a 96 well black bottom plates and treated with LLOME or Lipofectamine at varying concentrations. 24 hours after seeding, cells were stained with Hoechst 33342 (1 µg/mL) and digital images were collected and analyzed by an InCell 6000 by counting EGFP-Gal3 puncta/nuclei and averaged between 3 images. Cells with sgRNA knockdown were similarly analyzed using a comparable protocol 5 days after transduction with individual sgRNA-encoding lentivirus and puromycin selection of transduced cells.

### Cell fractionation and immunoblot

Cells were seeded into 3 mL at 250,000 cells/well in a 6-well dishes with 4.8 µl 0.43 mg/mL tau fibrils, and harvested after 48 hours by washing with PBS and releasing with 0.25% trypsin. Cells were resuspended with DMEM pre-warmed to 37 °C, spun down and washed again with PBS. Cells were resuspended in 20 µl PBS and lysed by flash freezing on dry ice and rapidly thawed at 42 C. This step was repeated twice. The resulting lysate was spun at 1000xg and the resulting supernatant was transferred to a new tube and respun to remove any carry-over insoluble material. The pellet was rinsed 3x with PBS and resuspended to the corresponding volume of supernatant and briefly sonicated with a tip sonicator (Sonopuls 2070) for a brief 1 second pulse at 10% maximum intensity. Equivalent fractions of total volume for 100 ng supernatant and resuspended pellet were boiled with SDS loading buffer (50 mM Tris-Cl (pH6.8), 2% (2 w/v) SDS, 0.1% (w/v) bromophenol blue) and 10 mM DTT, subjected to SDS-PAGE on 4-12% Bis-Tris polyacrylamide gels (ThermoFisher Cat# NP0322BOX) and transferred to nitrocellulose membranes. Primary antibodies against human tau (DAKO Cat# A0024) and β-actin (Cell Signaling Cat# 3700) were used to detect proteins. Blots were then incubated with secondary antibodies (Li-Cor Cat# 926-32213 and 926-68072) and imaged on the Odyssey Fc Imaging system (Li-Cor Cat# 2800). Digital images were processed and analyzed using Licor Image Studio™ software.

### qRT-PCR

CRISPRi-HEK293T cells expressing a constitutive non-targeting or targeting sgRNA were collected by centrifugation at 1000xg for 10 min, washed twice with ice cold PBS and processed for qPCR using a RNA purification kit (Zymo Cat# D7011). 500 ng total RNA from each sample were reverse transcribed using Superscript™ III reverse transcriptase using an oligo(dT) primer (Invitrogen Cat# 18080044). The resulting cDNA was diluted 5-fold using 10 mM Tris pH 8.0 and 0.67 µL of this dilution was used for each quantitative real-time PCR (qPCR) reaction. qPCR reactions were set up using SensiMix 2x Mastermix (Bioline Cat# QT615-20) and oligonucleotides targeting genes of interest (IDT) in triplicate and run on QuantStudio 6 Flex (Applied Biosystems Cat# 4485694) using protocols according to the mastermix manufacturer’s specifications. All reactions were normalized to an internal loading control (GAPDH) and the sgRNA activity is expressed as knockdown efficiency. The qPCR primer sequences are listed in Table S2.

## ACKNOWLEDGEMENTS

We thank Kathleen Keough and Nia Teerikorpi for contributing to preliminary studies. We thank all coauthors, Adam Frost, Bryce Mendelsohn, Jay Debnath, Amanda Woerman, Kartika Widjaja, Avi Samelson, Nina Dräger, Emmy Li, Poornima Ramkumar and other members of the Kampmann Lab for discussions and feedback on the manuscript. We thank Eric Chow and Derek Bogdanoff (UCSF Center for Advanced Technology) for support with next-generation sequencing and Sarah Elmes (UCSF Laboratory for Cell Analysis) for support with FACS. For human tissue samples, we thank Lea Grinberg, William Seeley and the Neurodegenerative Disease Brain Bank at the University of California, San Francisco.

## AUTHOR CONTRIBUTIONS

Conception and design: J.C.C., M.K.

Acquisition of data: J.C.C., D.L.N., P.R., M.N., E.T.

Analysis and interpretation of data: J.C.C., D.L.N., M.N., R.T., E.T., P.R., M.L., D.S., M.K. Drafting or revising the article: J.C.C., M.K.

Contributing unpublished essential data or reagents: J.Y.H., S.K.S., S.M., M.H., L.T.G., J.E.G.

## FOOTNOTES

The abbreviations used are

AD: Alzheimer’s Disease;
CRISPRi: CRISPR interference;
ESCRT: ndosomal sorting complexes required for transport;
FRET: fluorescence resonance energy transfer;
LLOME: Leucyl-Leucyl-o-Methyl-ester;
sgRNA: single guide RNA.

## Research support

This work was supported by NIH grants: New Innovator Award DP2 GM119139 (M.K.), R01 AG062359 (M.K.), R56 AG057528 (M.K. and L.T.G.), U54 NS100717 (M.K.), S10 OD021741 (supporting electron microscopy), R01 NS059690 (J.E.G.), P30 CA082103 (supporting the UCSF Laboratory for Cell Analysis), P01AG019724 and P50AG023501 (supporting the UCSF Neurodegenerative Disease Brain Bank), a Paul G. Allen Distinguished Investigator Award (M.K.), a Chan Zuckerberg Biohub Investigator Award (M.K.), the Tau Consortium (M.K., L.T.G., J.E.G. and the UCSF Neurodegenerative Disease Brain Bank), the Consortium for Frontotemporal Dementia Research (UCSF Neurodegenerative Disease Brain Bank), a QB3/Calico Longevity Postdoctoral Fellowship (J.J.C.), an Alzheimer’s Association Postdoctoral Fellowship (J.J.C.), a National Defense Science & Engineering Graduate Fellowship (S.K.S.), and an EMBO long-term postdoctoral fellowship ALTF 1193-2015 (M.Y.H.).

The content is solely the responsibility of the authors and does not necessarily represent the official views of the National Institutes of Health.

