## Supplementary material for "Compromised function of the ESCRT pathway promotes endolysosomal escape of tau seeds and propagation of tau aggregation": Figure S1

**Supporting Information:**
**Figure S1**

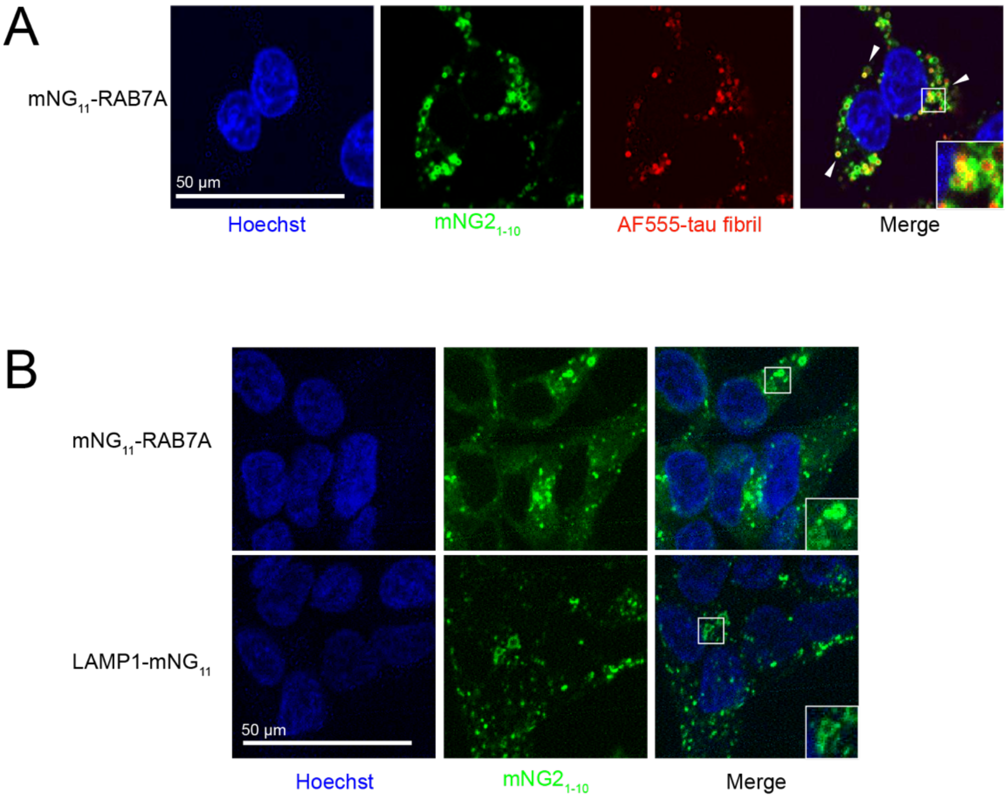


**Figure. S1 – Endogenous tagging of endolysosomal compartments**
(**A**) Representative fluorescence microscopy images of CRISPRi-HEK293T cells with RAB7A-mNG$_{11}$ endogenously labeled with the split-mNeonGreen system. Cells were treated with AF555-tau fibrils for 22 hours. (**B**) Representative fluorescence microscopy images of HEK293T cells with mNG$_{11}$-RAB7A (top) or LAMP1-mNG$_{11}$ (bottom) endogenously labeled with the split-mNeonGreen system.
